# Social facilitation of risky habitats in woodland caribou: responses to fire and roads

**DOI:** 10.1101/2024.10.01.616033

**Authors:** Jack G. Hendrix, Carissa D. Brown, John Gosse, Eric Vander Wal

## Abstract

Habitat change and subsequent trophic effects remain the dominant hypothesis for woodland caribou (*Rangifer tarandus*) population declines. Boreal forests are undergoing rapid changes, including expanding linear features and climate-induced increased wildfire activity. Within these landscapes caribou habitat selection and movement ought to maximize benefits while mitigating costs, e.g. predation risk, but spatial locations are likely not fixed in their cost-benefit ratio. For caribou, costs and benefits may vary seasonally and by social context. To test the effects of season and social context on caribou movement and selection of potentially risky habitats, we used a socially informed integrated step selection analysis (iSSA). We tested responses to two forms of risk in Terra Nova National Park, NL: roads and areas burned 2 – 75 years prior. Caribou avoided areas near roads irrespective of season. Burned area avoidance varied by season and burn age. Caribou avoided roads less and selected burns more in a social context. Social facilitation may permit the use of areas that were functionally unavailable to animals that were not in groups. As the most at-risk populations of woodland caribou exist at low density in highly disturbed landscapes, our results highlight the importance of considering sociality in maintaining access to resources.

## Introduction

Woodland caribou (*Rangifer tarandus*; hereafter caribou) populations are in decline throughout their range. Functional habitat loss and its subsequent effect on trophic interactions is the leading hypothesis for population declines (Festa-Bianchet et al. 2011). Caribou tend to avoid landscape disturbances such as linear features (Dyer et al. 2001), roads (Wilson et al. 2016), forestry (Courtois et al. 2008) or wildfire (Joly et al. 2003). In many cases habitat disturbance – be it forestry or fire – creates more open or early seral habitats, in which other ungulates appear to outcompete caribou and support increased predator populations, driving down caribou numbers (Latham et al. 2011). Unlike open habitats such as peatland complexes (Rettie and Messier 2000), open habitats generated by human disturbance or fire appear to be perceived as risky by caribou, leading to their selection for forage-poor mature forest stands and subsequent spatial separation from other ungulates and predators that tend to concentrate in early seral habitats (Cumming et al. 1996). Caribou that persist in disturbed landscapes are assumed to select habitat and move in a manner that maximizes their benefits over the costs. However, benefits and costs are not static in space and time. For example, costs and benefits vary by season (Leblond et al. 2011) and also by social context (Webber et al. 2024a). Here, we test how social context can facilitate selection of purportedly risky areas, i.e., near roads and previously burned areas, and to what extent this may vary seasonally.

Caribou evolved in landscapes that experience fire (Lafontaine et al. 2019). Fire regimes differ throughout their range, with fire return intervals in western Canada on the order of 100- 200 years vs. centuries to thousands of years for boreal forests in eastern Canada (Coops et al. 2018). Fires in eastern Canada are also smaller on average, with 5 – 10 km^2^ fires making up most of the area burned, in contrast with 200 – 500 km^2^ fires in western regions (Coops et al. 2018).

Anthropogenic climate change, however, is leading to increases in fire frequency and intensity (Kirchmeier-Young et al. 2019). Evidence from western Canada suggests caribou generally avoid recently burned areas (but see Mumma et al. 2018), though anthropogenically cleared areas, e.g., forestry cutblocks, may be avoided more strongly (SK: Skatter et al. 2017; YT: Russell 2018; AB: Stewart et al. 2020, Konkolics et al. 2021; but see BC: Mumma et al. 2018). These responses can vary seasonally, with some evidence of selecting burns during calving (Dekelaita et al. 2022) while avoiding burns over winter (Silva et al. 2020). Caribou avoid burned areas less as forests regenerate; 40 years is a commonly cited threshold (DeMars et al. 2019). However, fire cycles across the broad biogeographic range of caribou vary between ecoregions, thus 40 years is unlikely to apply universally. Time since fire may be particularly inapplicable in cases of failed regeneration, where post-disturbance coniferous forests convert to savannah or heathland rather than returning to their initial state (Mallik 1995). Failure to follow historical successional strategies is expected to occur more often with ongoing climate change (Stevens-Rumann et al. 2022) inducing more frequent fires (Brown and Johnstone 2012). Such an alternative successional pathway may result in more open habitats and extend the persistence of elevated risk in these burned areas, potentially indefinitely.

In addition to disturbances that convert habitat patches from one state to another, caribou also avoid linear features such as seismic lines or roads that fragment intact habitat (Dyer et al. 2002). Roads and other anthropogenic linear features functionally restrict caribou movement and act as semi-permeable barriers that caribou are reluctant to cross (Wilson et al. 2016). More heavily used roads have stronger effects that than those with less frequent human activity (Leblond et al. 2013; Smith and Johnson 2023). Aside from anthropogenic stressors and risk of direct collisions on roads (Leblond et al. 2013), linear features increase predator search efficiency and subsequently reduce both adult survival and reproductive success (Dickie et al. 2017; DeMars and Boutin 2018). While the physical footprint of such features is relatively small, caribou avoidance of roads leads to functional habitat loss of up to 1km in areas adjacent to roads (Schindler et al. 2007).

Caribou are unlikely to perceive risk of disturbed areas as static in geographic space. A given spatial location likely varies in risk (and reward) through time. The relative risks and rewards at a given location can change from an hourly scale (Lesmerises et al. 2018a) to seasonally (Hornseth and Rempel 2016). In seasonal environments, for example, availability of forage and the presence or absence of snow can drastically change the relative risks and rewards offered by certain habitats (Briand et al. 2009; Viana et al. 2018). Scarcity of winter forage encourages herbivores to take on more risk to find sufficient food, but deeper snow in open areas increases locomotory costs and mortality risk (Johnson et al. 2004; Eacker et al. 2023). Calves are far more vulnerable to predators than adult caribou, so perception of risk is amplified for females with calves-at-heel. In seasonally reproductive species where juvenile care and vulnerability is elevated in spring and summer, this can result in seemingly seasonal variation that is in fact driven by the presence of offspring (Leblond et al. 2011; Leclerc et al. 2012; Viejou et al. 2018).

Spatial and temporal variation in risk therefore creates a dynamic environment in response to which animals must adjust their spatial and social behaviours to balance costs and benefits. Social context that occurs in the same geographical space may also affect the ratio of risks to rewards (Webber and Vander Wal 2018; He et al. 2019; Webber et al. 2024b). Social associations in caribou and other ungulates influence movement and habitat selection across the landscape (Merkle et al. 2024; Webber et al. 2024a), with social influences on both the strength of selection for certain habitats and movement rates (Webber et al. 2024a). In fission-fusion systems where social interactions are flexible (Lesmerises et al. 2018b), group sizes are expected to increase with predation risk as group members benefit via increased collective vigilance (Creel et al. 2014) or dilution of risk (Mooring et al. 2004; Le Goff et al. 2024). In addition to selecting for shared locations with conspecifics, predation risk can also lead to increased sociality via movement synchrony among prey individuals (Prokopenko et al. 2024). Habitat is defined as the sum of resources, risks, and conditions (Matthiopoulos et al. 2023), but social context, e.g., group size, can also mediate risk through dilution and diminish rewards through increased competition for forage (Uccheddu et al. 2015). Social influences on behaviour are widespread even in seemingly asocial species (Makuya and Schradin 2024; Heeres et al. 2024). Thus, the importance of social interactions should not be discounted in spatial behaviour regardless of gregariousness.

We contrast two landscape features known to be perceived as risky to caribou to examine movement-integrated habitat selection and how seasonality and social context may mediate or attenuate perceived risk. Adopting the framework from (Dickie et al. 2020), we predict that animals will avoid, or select less, and move faster in risky habitats. Thus:

- We predict that caribou avoid roads, with higher-traffic roads avoided more (P1).
- We predict that caribou avoid areas in proximity to previously burned areas (P2), with greater avoidance of areas burned in the last 30 years.
- We predict that movement rates will increase in proximity to roads and previously burned areas (P3).
- Seasonally, we predict that female caribou during calving will be more risk averse than during the non-reproductive season. Thus, caribou during calving will more strongly avoid areas near roads and burns (P4).
- We predict that caribou in a more social context, i.e., in dyads, will be less averse to risky habitats. Thus, caribou will avoid less or select for areas near roads and burns (P5).

## Methods

### Caribou data

Adult female caribou were captured and outfitted with a Litetrack Iridium GPS collar (1250g, Lotek Wireless Inc., Newmarket, Ontario) from 9-10 March 2020. Caribou were immobilized by darting from helicopter using a mixture of Telazol and xylazine (1.5 mg/kg Telazol + 0.75 mg/kg xylazine) and were delivered a reversal agent to minimize total restraint time. Animal captures were conducted by Parks Canada following federal guidelines. Collars were programmed to record locations every 2 hours. Our initial dataset included GPS collar data for n = 10 individuals over a median of 656 days (1.8 years; min = 217 days, max = 979). Using these locations, we calculated straight line displacements (hereafter steps) between subsequent points, which formed the basis of our iSSA. We included only those steps occurring between consecutive 2-hour fixes and did not infer over missing observations or breaks in the GPS data. This resulted in a total of n = 10216 steps from n = 40040 point locations across all individuals. These steps were characterized by two main parameters: step length, or the distance between subsequent points analogous to speed, and turning angle relative to the prior step (where 0 = linear movement and 180 = complete reversal of direction). We took the set of steps for each individual and generated a distribution for step length and turning angles, then randomly sampled n = 10 theoretical available steps from this distribution to pair with each observed step. By comparing the end points of the observed step and its associated random steps, we can test whether caribou selected for or avoided certain conditions relative to their local availability.

To determine whether caribou spatial behaviour differed throughout the year, we separated our data into biological seasons (sensu Rudolph & Drapeau, 2012), following date ranges previously used to define seasonal home ranges for this population (J. Gosse, unpublished data.). To improve statistical power for seasons in which we had fewer steps, we pooled some periods to identify four biological seasons: winter (1 November – 31 March), spring migration (1 April – 19 May), calving and post-calving (20 May – August 30), and autumn (1 September – 31 October).

### Spatial and landscape data

We characterized habitat within our study area using the National Terrestrial Ecosystem Monitoring System’s 2020 land cover data (Hermosilla et al. 2022). This product uses Landsat imagery composites at 30 x 30m resolution to categorize pixels into one of 13 land cover classes. We simplified this system for our predominantly coniferous forested study area, pooling coniferous, broad leaf and mixedwood forest pixels as a single forest category. We also combined exposed/barren land, rock/rubble, and herbs as an open category. The remaining major land cover types – scrub, wetland, and water – were unchanged.

Fires and roads were our disturbances of interest. We used recent fire history records for the Terra Nova National Park (TNNP) region that ranged from 1945 to 2018 (Parks Canada Agency n.d.). Several fires in this area experienced regeneration failure and converted to an open Kalmia heath ecosystem (*Kalmia angustifolia*; Mallik et al. 2010). Based on expert opinion from local land managers and a natural breakpoint in the chronosequence, we chose to separate fires into ‘younger’ (post-1992, < 30 years post-fire) and ‘older’ (pre-1992, > 30 years post-fire).

Burn polygons occupied a small area (2477 ha) of our study area (198,328 ha for 99% kernel density, burns = 1.2%), and relatively few caribou locations occurred within burns themselves. We thus elected to use the distance to the nearest older and younger burns, rather than considering burns as a land cover type themselves. We assigned a distance of 0 to points within burns. Using distance to burn at the end point of each observed and available step, we could determine whether caribou were selecting to be nearer or farther from younger and older burns.

We similarly calculated the distance from each end point to the nearest major and minor roads within the study area using OpenStreetMap data (Padgham et al. 2017). The major road in this region is the high-use Trans Canada Highway (TCH), which roughly bisects Terra Nova National Park running north-south. The remaining roads in the area are minor access roads with low traffic volumes. We extracted distances to the TCH and minor access road at the end point of each step as described above.

### Model structure

We designed two parallel models, one to test for caribou response to roads and the other for the response of caribou to burned areas, as models including both consistently failed to converge. Distance to major and minor roads and distance to older and younger burns were log transformed in the model as we expected the strength of response to exponentially decay with increasing distance. In addition to our primary disturbances of interest, both models also included landcover at the end of each step. Here, land cover was simplified as two dichotomous variables: forest vs. not forest and open vs. not open. We included all these spatial covariates to test for caribou selection or avoidance of each variable. We also interacted each with step length, which enabled us to identify whether caribou travelled faster or slower in specific habitats or in proximity to burns or roads. Each covariate and covariate-step length interaction were also interacted with individual ID so that we could differentiate individual responses to these landscape features. All model structures are provided in the Appendix.

We ran the road model and fire model on the full dataset, and subsequently ran the same model structure separately on each of our four biological seasons. We compared the effects of each predictor in a given season to its respective annual model to determine how caribou spatial behaviour and responses varied throughout the year. Finally, we were interested in how sociality would mediate spatial behaviour in this population. To remotely determine sociality, we used the gambit of the group rule where GPS fixes from two collared individuals within a 50m radius at the same time indicate dyad formation between these individuals (Robitaille et al. 2018). While only a subset of the population was collared, our sample of 10 individuals represented 17% of females in the local population. Though we cannot fully determine whether caribou were truly solitary when other collared animals were not nearby, this metric represents a gradient of sociality. Gregariousness in this population is relatively low throughout the year, particularly during the calving period as in other Newfoundland caribou herds (Hendrix et al. 2024). The winter season was the only period in which we had sufficient dyads for statistical comparison (17% of steps occurred in a dyad vs. < 6% in other seasons). We thus restricted our ‘social’ models to only this season. We used a dichotomous measure of in dyad vs. not in dyad at the beginning of each step, and interacted this sociality term with the land cover and distance-to predictors described above, to develop a ‘social road’ and ‘social fire’ model using the winter seasonal data. Using these models, we were able to ascertain whether being in a dyad mediated caribou habitat selection and how they responded to risky features on the landscape. We were unable to include three-way interactions of dyad with step length and other spatial predictors, as this proved too complex a model for the existing data (i.e., convergence was not achieved). As such we did not explicitly test how sociality affected movement rates in this system.

All analyses were conducted in R version 4.3.2 (R Core Team, 2023), using the amt (Signer et al. 2019), glmmTMB (Brooks et al. 2017), sf (Pebesma 2018), spatsoc (Robitaille et al. 2019), and targets (Landau 2021) packages.

## Results

In all models, caribou selected for forest over open habitats but moved faster in open areas when they were used (Appendix, Fig S1). These effects persisted across seasons and when both roads or burned areas were included as covariates.

### P1 – response to roads

Caribou avoided locations near the Trans-Canada Highway (TCH) and near minor roads (Fig 1). Responses to minor roads were highly conserved across individuals, while there was some variation in response to the TCH, with one caribou exhibiting the opposite trend and selecting to be nearer the highway.

**Figure 1.**
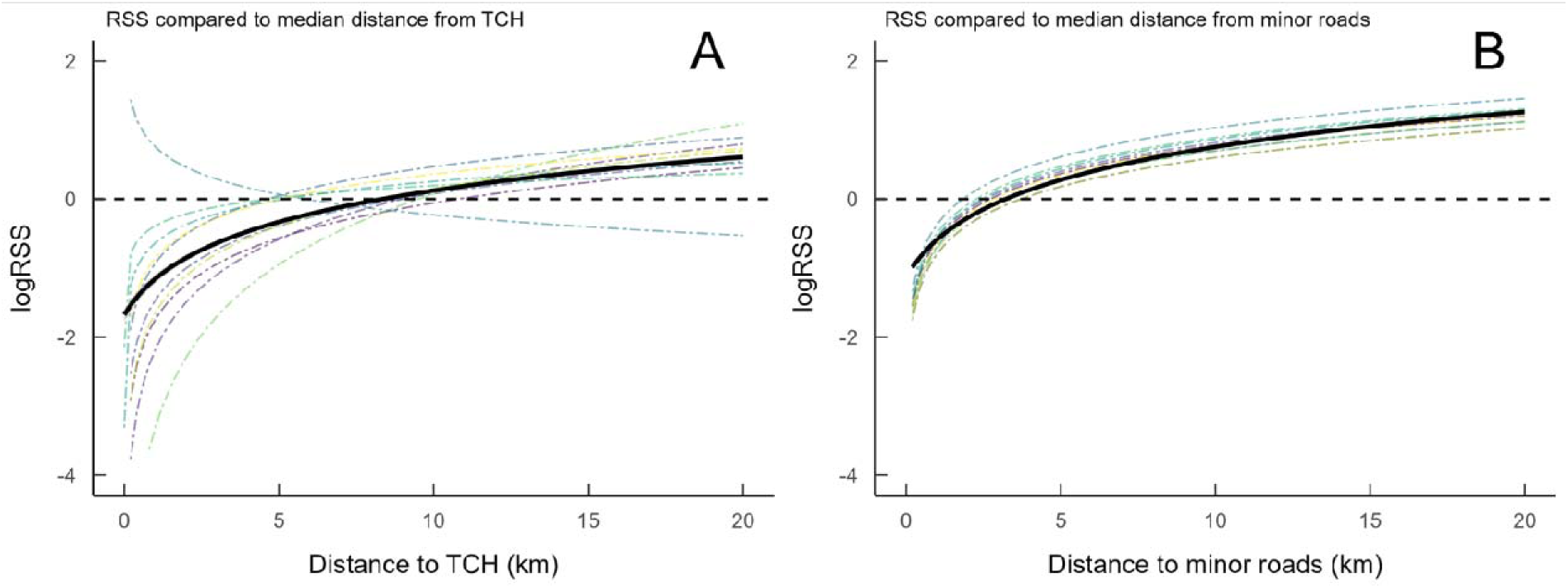
Relative selection strength (RSS) in response to distance to roads for caribou in TNNP. Each coloured dashed line represents an individual caribou, with the solid black line showing the population average. RSS is the strength of selection for a given value of a covariate, compared to a reference level. In this case values between 0 and 20 km from roads are compared against the median distance for each individual. The y-axis is log() transformed, so a y-value of zero represents a relative selection strength of 1, or a neutral response (thus lines cross the x-axis at the median value). Values less than 0 reflect avoidance, and values greater than zero reflect selection. (A) Most caribou were more likely to select points further from the Trans-Canada Highway (TCH), i.e. caribou avoided areas near the highway, although one individual showed the opposite response. (B) Caribou were more likely to select points further from minor roads, i.e. caribou avoided areas near minor roads.

### P2 – response to fires

Caribou responses to burns were more individually variable than responses to roads. Caribou avoided locations near younger (< 30 year old) burns at the population level, though two animals exhibited the opposite effect by selecting proximity to younger burns (Fig 2A). In contrast, the population on average selected for proximity to older (> 30 year old) burns, although again there were two individuals that exhibited the opposite response (Fig 2B). The magnitude of response to both older and younger burns was substantially smaller than the response to either type of road.

**Figure 2.**
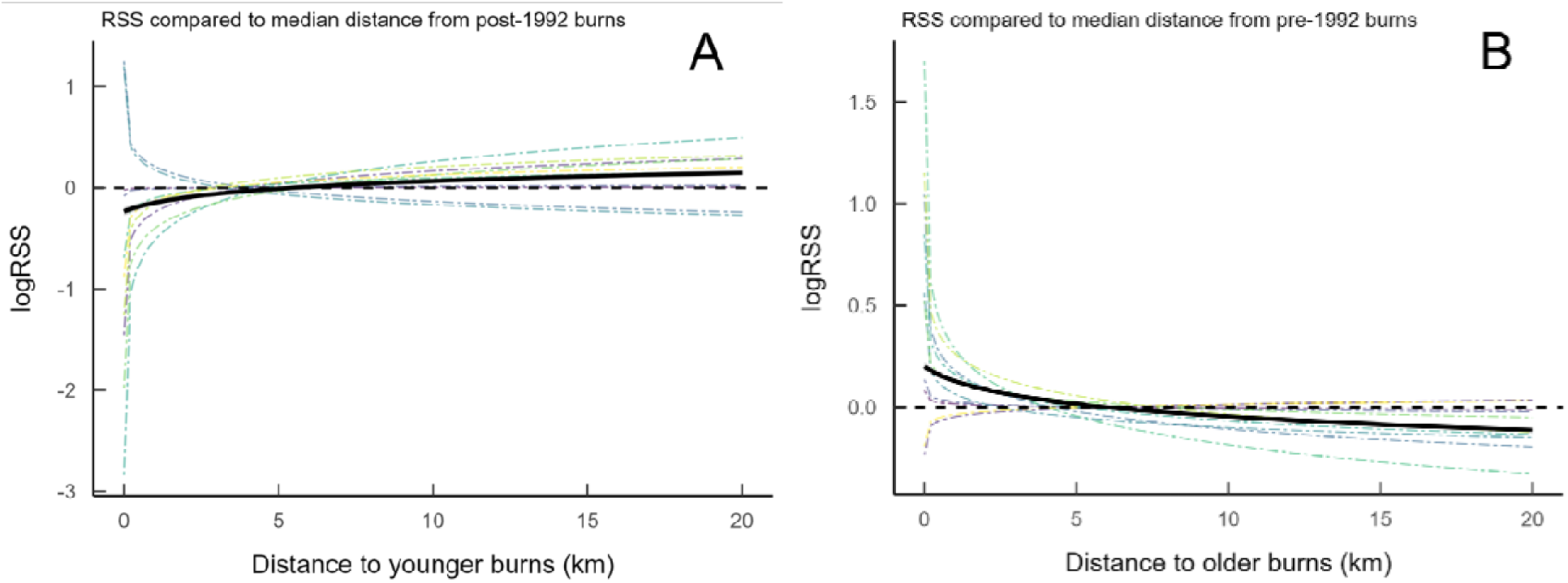
Relative selection strength (RSS) in response to distance to past fires for caribou in TNNP. Each coloured dashed line represents an individual caribou, with the solid black line showing the population average. RSS is the strength of selection for a given value of a covariate, compared to a reference level. In this case values between 0 and 20 km from are compared against the median distance for each individual (approx. 5 km). The y-axis is log() transformed, so a y-value of zero represents a relative selection strength of 1, or a neutral response (thus lines cross the x-axis at the median value). Values less than 0 reflect avoidance, and values greater than zero reflect selection. (A) Most caribou were less likely to select points closer to younger burns, i.e., caribou avoided areas near younger burns, although two individuals showed an opposite response. (B) Most caribou were more likely to select points closer to younger burns, i.e. caribou selected areas near older burns, although two individuals showed an opposite response where they were more likely to avoid locations near younger burns.

### P3 – Movement rate near risk

Caribou movements increased most with proximity to the TCH. Movement rate also increased near minor roads, in line with our prediction. Caribou moved faster near older burns but movement was unchanged in proximity to younger burns (Fig 3). This contradicts our prediction that movement rate should increase in riskier areas, particularly since younger burns were avoided more intensely than older burns. There was also a wide range of individual variation in movement rates, with some caribou consistently moving faster than others. The difference between the fastest and slowest individuals was larger than the effect of any spatial predictor on movement rate.

**Figure 3.**
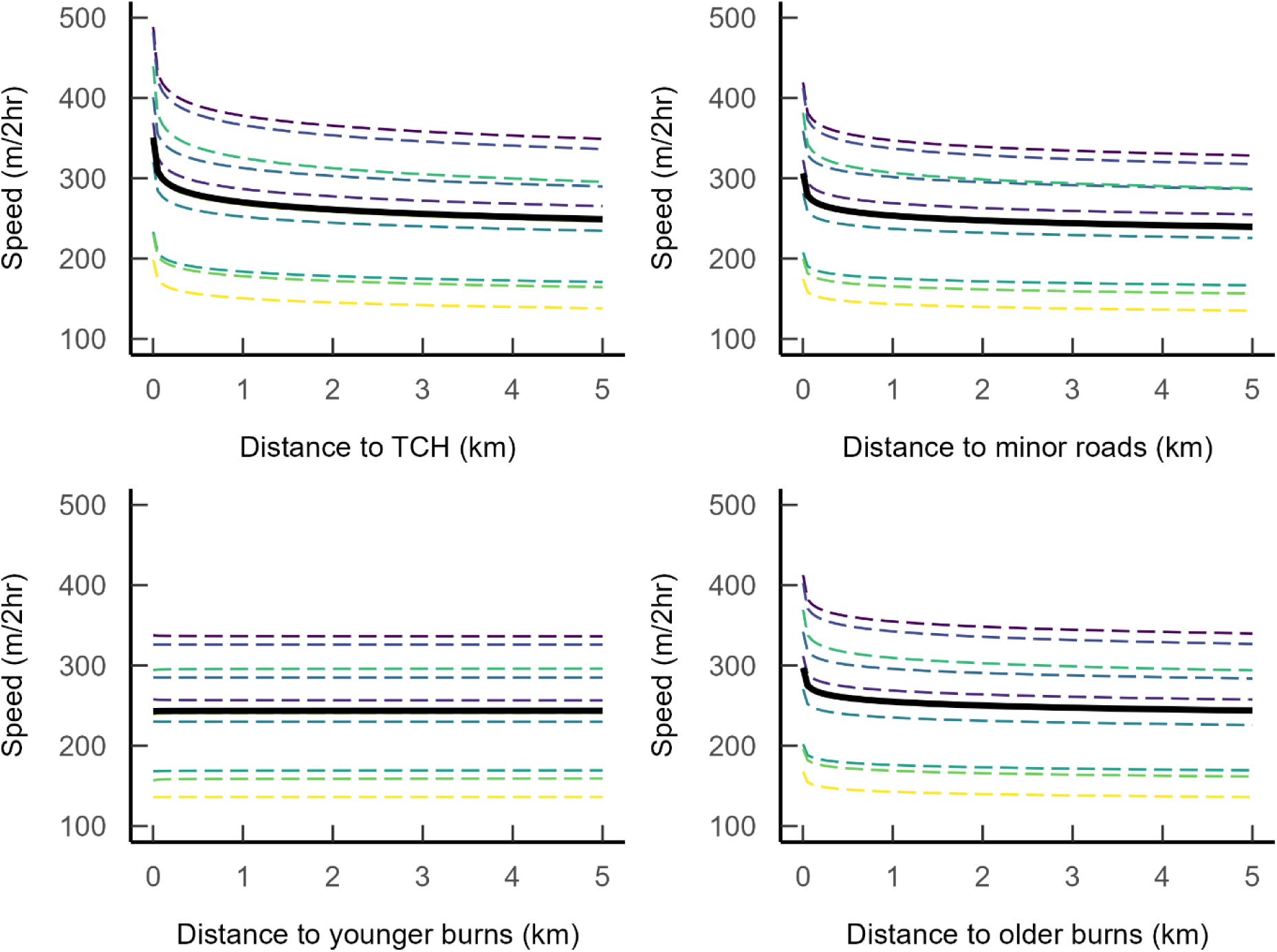
Changes in movement rate of woodland caribou based on distance to various landscape features. Each coloured dashed line represents an individual caribou, with the solid black line showing the population average effect.

### P4 – Seasonality of risk

Caribou avoidance of both younger and older burns was strongest during the calving and post-calving season (20 May – 30 August), which was consistent with our prediction that caribou would be most risk averse during these important biological seasons (Figs 4, 5). The response of caribou to both younger and older burns was essentially neutral during winter (1 November – 31 March). This trend was also observed for younger burns during autumn (1 September – 31 October), while older burns were selected during autumn. During spring migration (1 April – 19 May) younger burns were generally avoided. There was dramatic individual difference in response to older burns during spring migration: the population average response was neutral, but this is the result of strong and opposing selection and avoidance of older burns by various individuals cancelling out.

**Figure 4.**
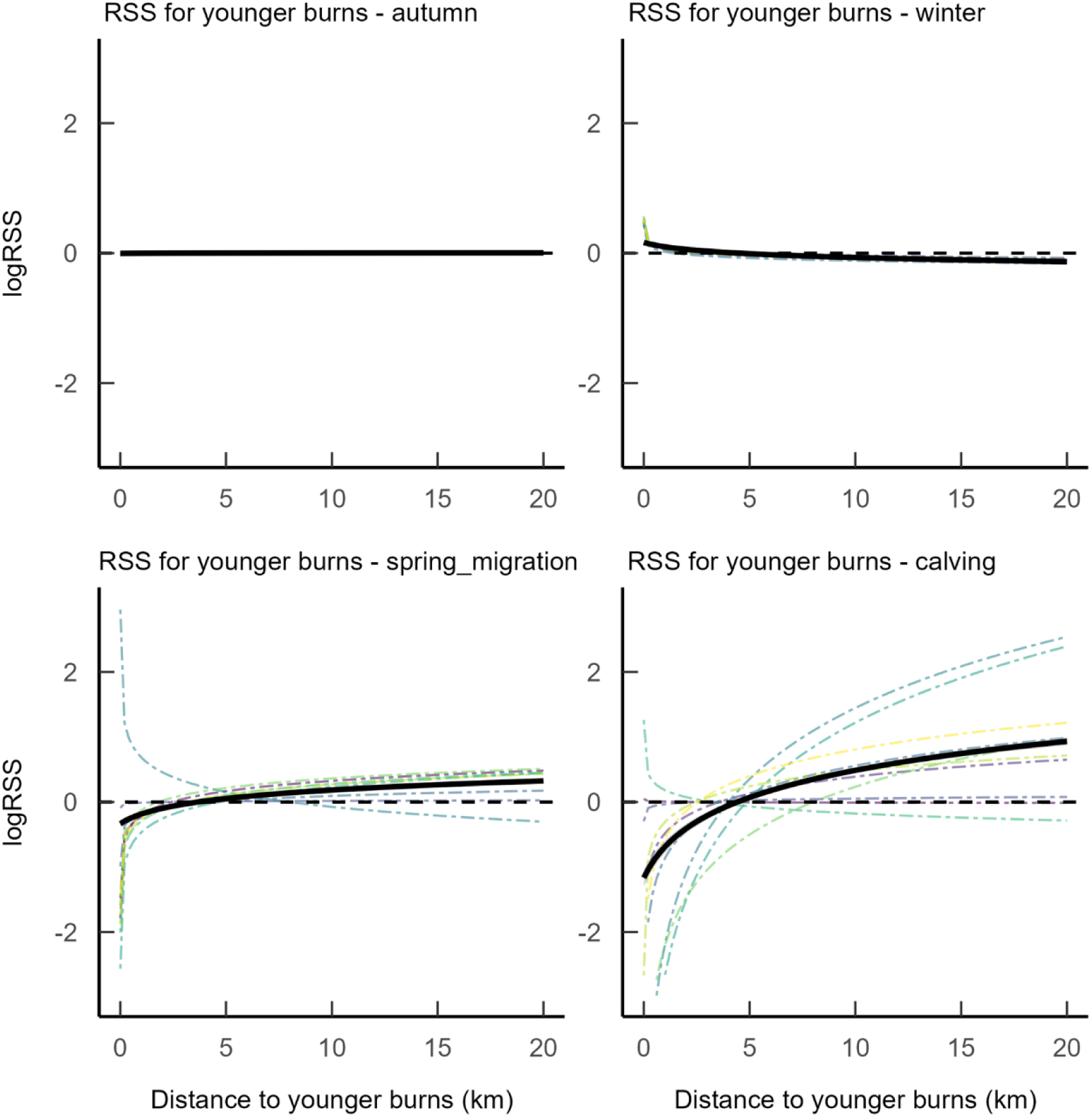
Seasonal differences in relative selection strength (RSS) in response to distance to younger burns for Terra Nova caribou. Each coloured dashed line represents an individual caribou, with the solid black line showing the population average. RSS is the strength of selection for a given value of a covariate, compared to a reference level. In this case values between 0 and 20 km from are compared against the median distance for each individual (approx. 5 km). The y-axis is log() transformed, so a y-value of zero represents a relative selection strength of 1, or a neutral response (thus lines cross the x-axis at the median value). Values less than 0 reflect avoidance, and values greater than zero reflect selection.

**Figure 5.**
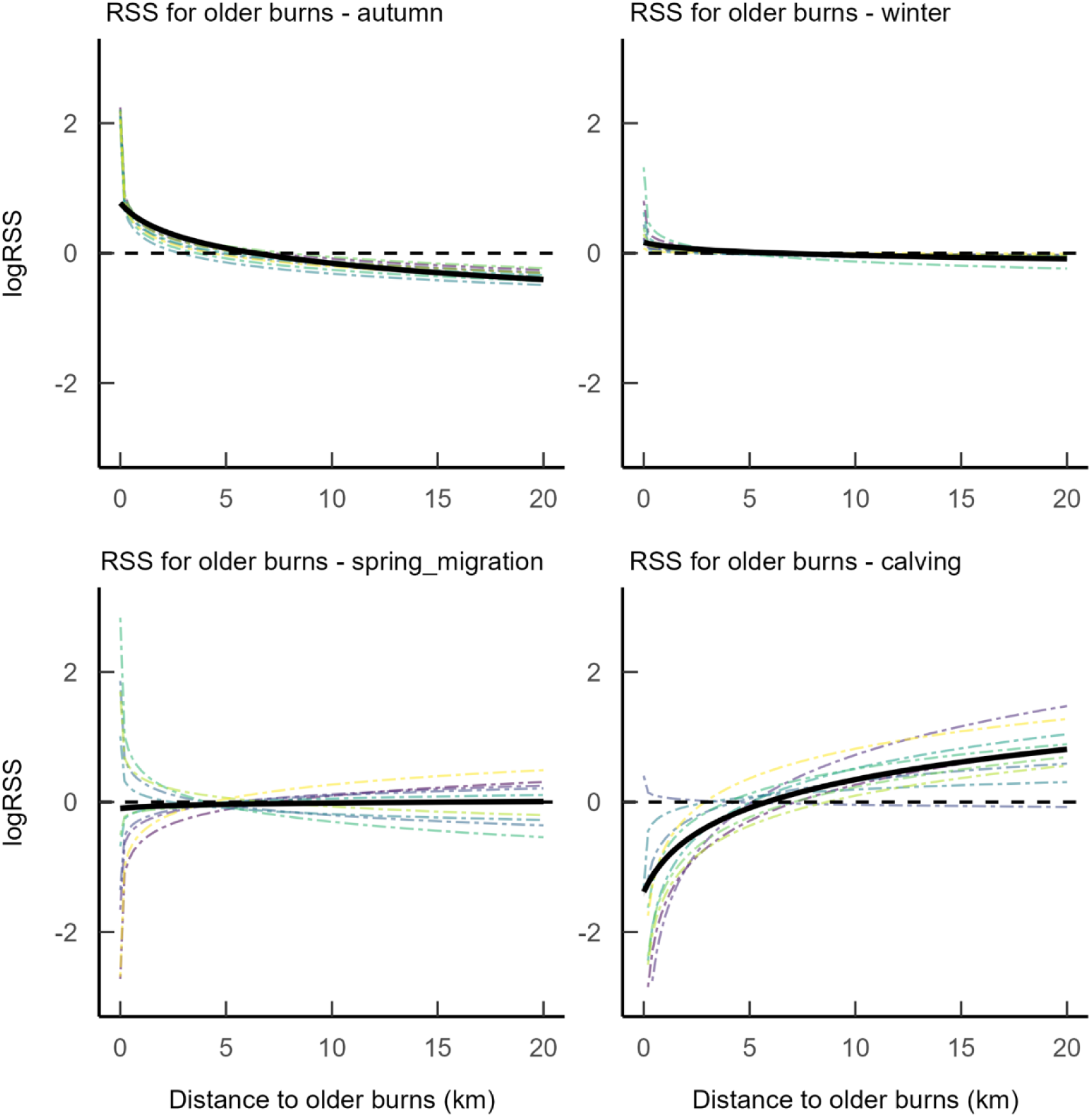
Seasonal differences in relative selection strength (RSS) in response to distance to older burns for Terra Nova caribou. Each coloured dashed line represents an individual caribou, with the solid black line showing the population average. RSS is the strength of selection for a given value of a covariate, compared to a reference level. In this case values between 0 and 20 km from are compared against the median distance for each individual (approx. 5 km). The y-axis is log() transformed, so a y-value of zero represents a relative selection strength of 1, or a neutral response (thus lines cross the x-axis at the median value). Values less than 0 reflect avoidance, and values greater than zero reflect selection.

Caribou responses to the TCH were less seasonally variable than responses to burns, however there was pronounced avoidance of the highway during the winter season (Fig 6). The average response of the population to the TCH during calving was weak avoidance, but this average obscures stronger individual responses of some caribou selecting and others avoiding the highway during this time. In contrast, minor roads, were consistently avoided during all seasons with only minimal differences in the strength of this relationship (Fig S2).

**Figure 6.**
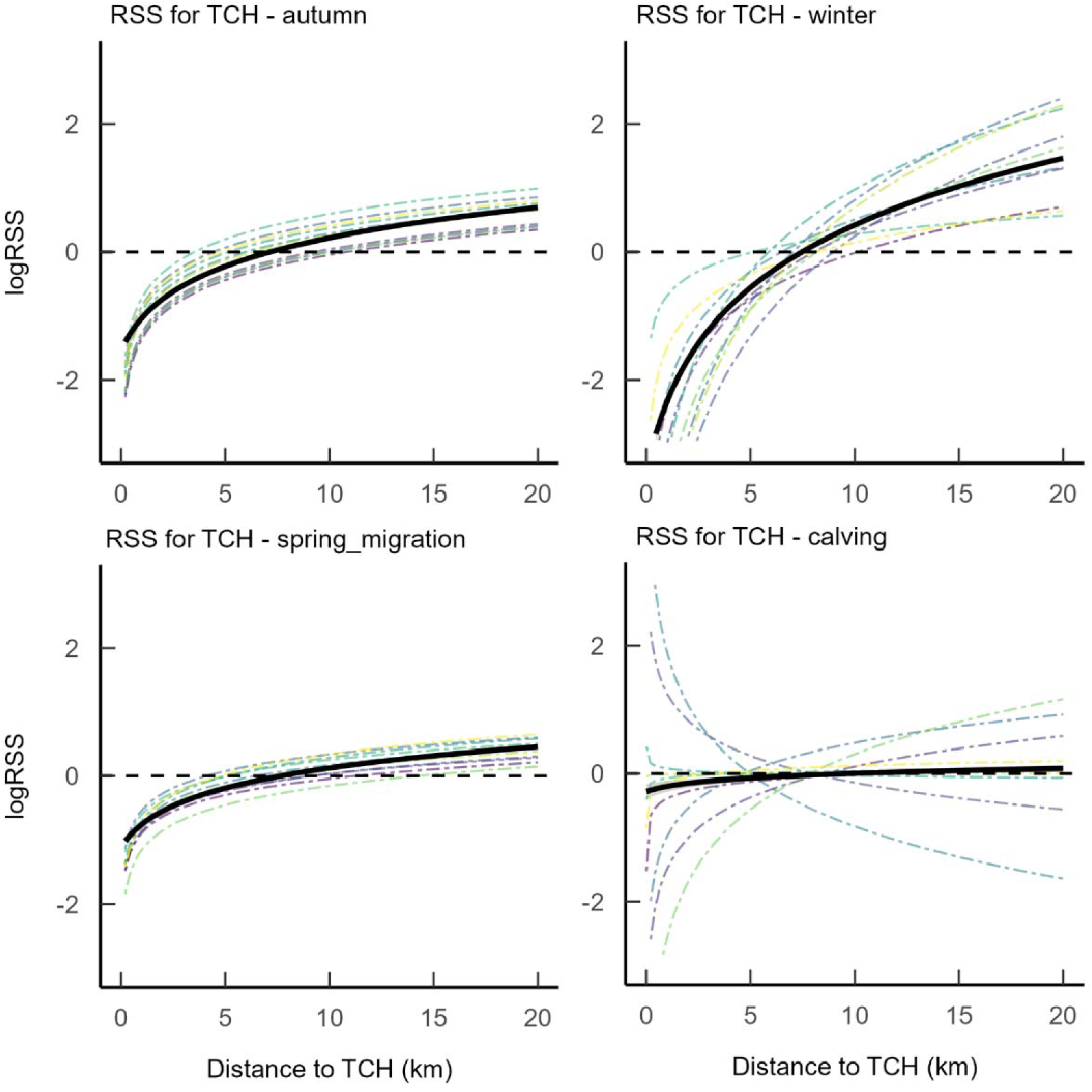
Seasonal differences in relative selection strength (RSS) in response to distance to the Trans Canada Highway (TCH) for Terra Nova caribou. Each coloured dashed line represents an individual caribou, with the solid black line showing the population average. RSS is the strength of selection for a given value of a covariate, compared to a reference level. In this case values between 0 and 20 km from are compared against the median distance for each individual (approx. 8 km). The y-axis is log() transformed, so a y-value of zero represents a relative selection strength of 1, or a neutral response (thus lines cross the x-axis at the median value). Values less than 0 reflect avoidance, and values greater than zero reflect selection.

### P5 – Social mediation of risk

Social context mediated selection for several habitat features. Caribou selected forests in all our asocial models, but when social context was included, we found that caribou in dyads actually avoided forested areas during winter while the selection for forest was driven by caribou not in dyads (Fig S3). Caribou in dyads did not avoid roads as intensely as caribou not in dyads (Fig 7), suggesting the presence of conspecifics made individuals less risk averse. Caribou selected more strongly for both younger and older burns during the winter season when in dyads. This social effect was particularly strong for younger burns (Fig 8).

**Figure 7.**
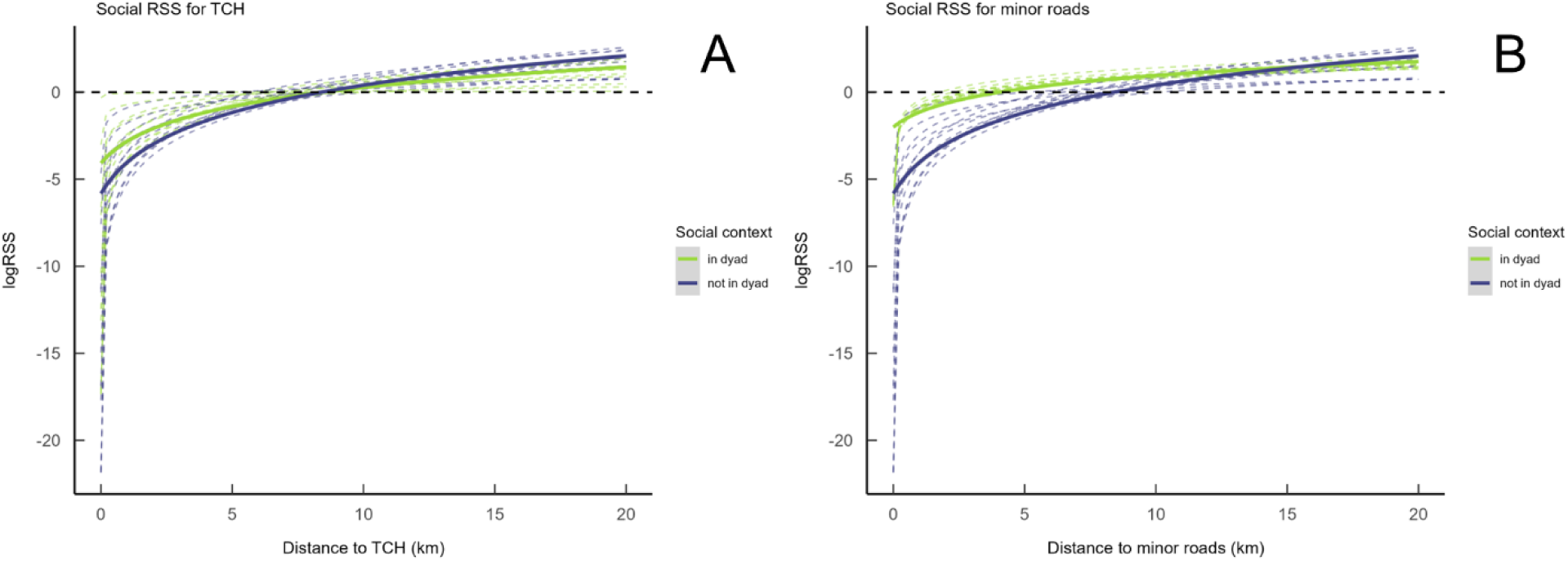
Relative selection strength (RSS) for distance to Trans-Canada Highway (TCH; A) and minor roads (B) by caribou during winter. Green lines represent periods when caribou were in a dyad with other collared individuals. Blue lines are periods when no other collared individuals were within 50m of the focal animal. RSS is the strength of selection for a given value of a covariate, compared to a reference level. In this case values between 0 and 20 km from are compared against the median distance for each individual (approx. 8 km). The y-axis is log() transformed, so a y-value of zero represents a relative selection strength of 1, or a neutral response (thus lines cross the x-axis at the median value). Values less than 0 reflect avoidance, and values greater than zero reflect selection.

**Figure 8.**
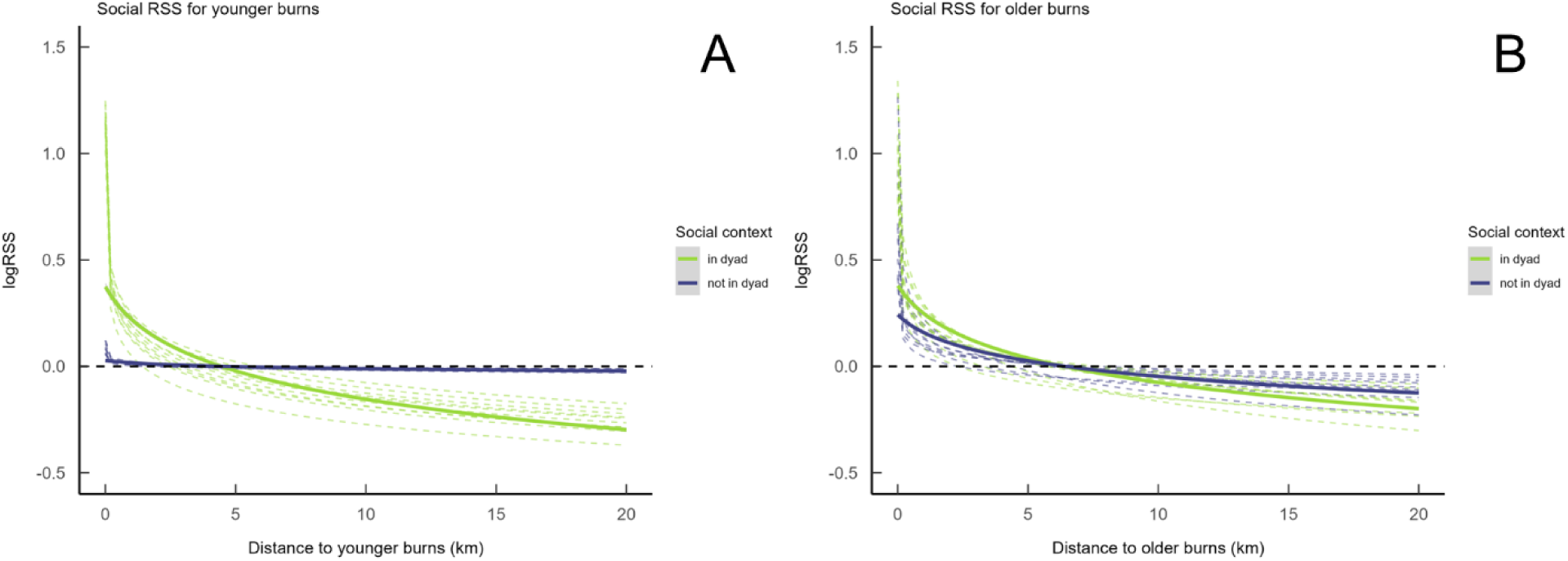
Relative selection strength (RSS) for distance to younger (A) and older (B) burns by caribou during winter. Green lines represent periods when caribou were in a dyad with other collared individuals. Blue lines are periods when no other collared individuals were within 50m of the focal animal. RSS is the strength of selection for a given value of a covariate, compared to a reference level. In this case values between 0 and 20 km from are compared against the median distance for each individual (approx. 5 km). The y-axis is log() transformed, so a y-value of zero represents a relative selection strength of 1, or a neutral response (thus lines cross the x-axis at the median value). Values less than 0 reflect avoidance, and values greater than zero reflect selection.

## Discussion

How woodland caribou use the landscapes in which they persist is critical in the management of at-risk populations. Conservation measures generally strive to protect locations identified as key habitat that are consistently selected by caribou (COSEWIC 2014). While it is well-established that landscape disturbances like roads and burns have negative effects on caribou and these effects can vary by season, we found that social context mediated habitat selection of such risky places. In TNNP, caribou responses to burned areas were more seasonally variable than responses to roads. Caribou consistently perceived roads as risky and avoided them across seasons. In both cases, social context changed how caribou associated with roads and previously burned areas. Caribou were less risk averse when other collared conspecifics were nearby.

Caribou avoided roads less and select for previously burned areas more during the winter season.

We predicted (P1) that caribou would avoid locations near roads, and avoid major roads to a greater extent than less trafficked roads. Indeed, caribou avoided the Trans-Canada Highway more strongly overall but were also more individually variable in their selection. Caribou responded to minor roads more consistently across individuals and seasons. Caribou also moved faster in proximity to roads, indicative of these areas being perceived as risky (P3; Matthiopoulos et al. 2023). The effects of linear features on caribou habitat selection and movement are well-documented (Serrouya et al. 2020; Walker et al. 2024), and the increased avoidance of larger and more intensely used linear features we observed is consistent with earlier studies. Notably, much of the linear feature research in western North American boreal caribou populations focuses on caribou avoiding roads and seismic lines because these function as movement corridors for wolves and increase predation risk (Dickie et al. 2017; DeMars and Boutin 2018). Wolves were extirpated from the island of Newfoundland nearly a century ago (Bergerud 1971). As coyotes (*Canis latrans*) and black bears (*Ursus americanus*) primarily predate upon caribou calves on insular Newfoundland (Mahoney et al. 2016), adult predation risk is unlikely to be a major driver of the avoidance behaviour observed in our system. Human activity on roads may be the more dominant driver of their perceived risk, rather than a predator-mediated movement effect.

Caribou were more likely to be harvested by hunters in Newfoundland when closer to roads and towns (McNamara et al. 2022). Although caribou hunting is not permitted within TNNP, this avoidance behaviour likely persists given the movement of animals across park boundaries to surrounding areas. While caribou likely perceived roads solely as sources of risk, caribou behaviour around previously burned areas was more nuanced.

We predicted (P2) that caribou avoid burned areas, with less avoidance of older burns, but caribou did not universally avoid previously burned areas in this system. Much of the research on caribou-fire interactions has focused on western portions of woodland caribou range (Rickbeil et al. 2017; Skatter et al. 2017; DeMars et al. 2019; Silva et al. 2020; but see Lafontaine et al. 2019). Western Canadian boreal forests, being both hotter and drier (Johnson et al. 1998), generally have a shorter fire return interval (i.e., less time between fires) and more severe fires than the eastern boreal (Hart and Chen 2006). Newfoundland in particular has a wet maritime climate, and caribou populations on the island likely have much less frequent exposure to wildfire (mean fire interval of 250 years for Terra Nova, Lake et al. 2024, with regional estimates >700 years in central Newfoundland forests, Arsenault et al. 2016) than caribou in British Columbia or Alberta where the bulk of caribou-fire research has occurred (Konkolics et al. 2021; Maltman et al. 2024). Western boreal caribou populations also face much higher levels of habitat disturbance overall within their range from forestry and resource extraction in addition to fire (Sorensen et al. 2008) than in Newfoundland where the human footprint remains comparatively restricted (McCarthy et al. 2011). Caribou from regions with more regular fire activity exhibit greater avoidance of recent burns, while those with less experience of wildfire were more likely to select disturbed habitats (Lafontaine et al. 2019). Central Newfoundland herds selected for 10- and 20-year-old burns during calving, and this was associated with improved calf recruitment (Dekelaita et al. 2022). The herds investigated by Dekelaita et al (2022), however, exhibit a different calving behaviour than the caribou in TNNP. The large migratory herds in central Newfoundland, e.g., Middle Ridge and La Poile, congregate in non-forested calving grounds during parturition (Rayl et al. 2014; but see Bonar et al. 2020, calving aggregations may be density-dependent). Selection for areas cleared by fire is an effective anti-predator strategy if the population relies on aggregation and group vigilance in open habitats. Conversely, caribou in TNNP and other smaller herds tend to disperse and give birth in isolation (Rayl et al. 2014; Hendrix et al. 2024), relying on dense forest to reduce predator detection; avoidance of burns and other open areas is a risk-minimizing strategy in this social context.

Consistent with the hypothesis that caribou are more risk averse during calving, we found that avoidance of burns peaked during calving season (P4). We did not find any such seasonal trends for caribou responses to roads. We submit that the open habitat created by burns may offer relative benefits to caribou during certain times of year, while roads offer no such rewards. We interpret selecting burns more during autumn and winter as likely related to their benefits, while avoidance peaking during calving spotlights the amplified risk. For instance, caribou forage is relatively scarce over winter compared to summer (Bergerud 1974). During winter, caribou diets shift to become lichen-dominated as fresh, more nutritious, vegetation becomes unavailable (Webber et al. 2022). While the Kalmia that generally dominates post-fire in Newfoundland is known to be consumed by caribou (Bergerud 1972), we propose it is likely incidental or accidental forage due to its very high tannin concentrations (Joanisse et al. 2007) that diminish its utility as nutritious forage. Indeed as the common names sheep laurel and lambkill suggest, Kalmia is toxic to livestock (Marsh and Clawson 1930). However, Kalmia heaths often support extensive *Cladonia* spp. lichen beds (Mallik and Kayes 2018) that may be particularly attractive as winter foraging patches (Bergerud 1974). The increased avoidance of burns during calving that we observed aligns with the prediction that females with offspring perceive these open areas as riskier, coupled with increased availability of non-lichen foraging opportunities elsewhere in the summer. Conversely, the absence of such seasonal variation in response to roads suggests that the risk-reward balance for these features is static. Thus, roads are likely perceived as more uniformly risky on the landscape, without associated ecological benefits.

Social groups form to reduce per-capita risk (Silk 2007). Thus, we predicted social facilitation would present as selection or reduced avoidance for risky landscape features when in the presence of conspecifics (P5). We attribute the seasonal changes in habitat selection that we observed to increased caribou gregariousness during winter. Many woodland caribou populations in Newfoundland are mostly solitary during calving and aggregate in winter (Webber and Vander Wal 2021; Le Goff et al. 2024) to make use of cues from conspecifics to find patchy resources (Peignier et al. 2019). Caribou in dyads were less avoidant of both types of roads than caribou not observed in dyads. Furthermore, when not in dyads caribou had an equivocal or neutral response to younger burns, but caribou in dyads selected these areas. Caribou in dyads also selected more strongly for older burns than caribou not in dyads. The potential lichen foraging opportunities in burned areas during winter were effectively accessible only to caribou in dyads, as individuals not in dyads had much weaker selection for these areas. We acknowledge that because we only had one in six adult females collared, we cannot demonstrate animals that were not in dyads were necessarily solitary, nor were we able to conclude that animals in dyads were not also in larger groups with uncollared individuals. Our dyadic metric, however, reflects relative sociality, and we detected consequences of this difference on habitat selection.

Functional habitat loss is contributing to woodland caribou declines in Canada are (Stewart et al. 2020; Nagy-Reis et al. 2021). Spatial responses to disturbance are therefore central to our management of woodland caribou. We tested how seasonality and sociality affected how caribou may have perceived two forms of risk implicated in functional habitat loss: roads and previously burned areas. Woodland caribou in and around Terra Nova National Park typically avoided areas near roads and near post-fire habitats. The response to burns, however, varied by season with caribou switching from avoidance to selection of burns in winter, particularly so when in a social context. Caribou associating with conspecifics also avoided roads less so than caribou not in dyads, effectively shrinking the zone of influence around these linear features. Conservation of caribou rarely considers social behaviour. If caribou can functionally access some habitats only in the presence of conspecifics, social facilitation could have implications for caribou conservation. For example, in small populations experiencing inverse density dependence, i.e., Allee effects, where population declines accelerate as density decreases (Wittmer et al. 2005), group formation and social interactions are likely to be increasingly rare as density falls (Courchamp et al. 1999). Therefore, small, sparsely distributed populations may have less functional, or accessible, habitat than larger populations. Woodland caribou populations have declined sharply and many herds persist at much lower density than historical numbers in an increasingly disturbed landscape (Festa-Bianchet et al. 2011). Currently observed habitat use in diminished populations might reflect only those areas available in the absence of social facilitation. If such populations recover to sufficient density for groups to form more often or at all, our results suggest caribou may make use of areas previously considered functionally unavailable.

## Acknowledgements

We respectfully acknowledge that the data collection and analysis for this project occurred in Ktaqmkuk, what is now known as the island of Newfoundland, the ancestral homelands of the Beothuk and Mi’kmaq peoples. We are very grateful to members of the Wildlife Evolutionary Ecology Lab for their feedback on early drafts, and to Christina Prokopenko, Alec Robitaille, and Julie Turner for their invaluable assistance in constructing and troubleshooting our iSSA models. We thank John Neville (Department of Environment and Conservation) and Chris Gosse (Newfoundland Helicopters Ltd.) for deploying GPS satellite collars.

## Author contributions

J.G.H.: Conceptualization, Formal Analysis, Methodology, Writing – original draft, Writing – review & editing, Visualization. C.D.B.: Writing – review & editing. J.G.: Conceptualization, Funding Acquisition, Resources, Writing – review & editing. E.V.W.: Supervision, Writing – review & editing.

## Data availability statement

We have made a version of our data available, along with code to replicate analyses, in a Github repository at https://github.com/jghendrix/social-risk-TerraNova-caribou. Due to sensitivity around sharing location data for this population, we have removed XY coordinates from this dataset, but retained the stepwise and landscape covariates required for our models included here.

## References

1. Arsenault, A., LeBlanc, R., Earle, E., Brooks, D., Clarke, B., Lavigne, D., and Royer, L. 2016. Unravelling the past to manage Newfoundland’s forests for the future. The Forestry Chronicle 92(04): 487–502. doi:10.5558/tfc2016-085.

2. Bergerud, A.T. 1971. The population dynamics of Newfoundland caribou. Wildlife Biology Monographs 25: 3–55. doi:10.2307/3830629.

3. Bergerud, A.T. 1972. Food habits of Newfoundland caribou. The Journal of Wildlife Management 36(3): 913–923. doi:10.2307/3799448.

4. Bergerud, A.T. 1974. Relative abundance of food in winter for Newfoundland caribou. Oikos 25(3): 379– 387.

5. Bonar, M., Lewis, K.P., Webber, Q.M.R., Dobbin, M., Laforge, M.P., and Vander Wal, E. 2020. Geometry of the ideal free distribution: individual behavioural variation and annual reproductive success in aggregations of a social ungulate. Ecology Letters 23: 1360–1369. doi:10.1111/ele.13563.

6. Briand, Y., Ouellet, J.-P., Dussault, C., and St-Laurent, M.-H. 2009. Fine-scale habitat selection by female forest-dwelling caribou in managed boreal forest: Empirical evidence of a seasonal shift between foraging opportunities and antipredator strategies. Écoscience 16(3): 330–340. doi:10.2980/16-3-3248.

7. Brooks, M.E., Kristensen, K., Benthem, K.J. van, Magnusson, A., Berg, C.W., Nielsen, A., Skaug, H.J., Mächler, M., and Bolker, B.M. 2017. glmmTMB Balances Speed and Flexibility Among Packages for Zero-inflated Generalized Linear Mixed Modeling. The R Journal 9(2): 378–400.

8. Brown, C.D., and Johnstone, J.F. 2012. Once burned, twice shy: Repeat fires reduce seed availability and alter substrate constraints on *Picea mariana* regeneration. Forest Ecology and Management 266: 34–41. doi:10.1016/j.foreco.2011.11.006.

9. Coops, N.C., Hermosilla, T., Wulder, M.A., White, J.C., and Bolton, D.K. 2018. A thirty year, fine-scale, characterization of area burned in Canadian forests shows evidence of regionally increasing trends in the last decade. PLOS ONE 13(5): e0197218. doi:10.1371/journal.pone.0197218.

10. COSEWIC. 2014. COSEWIC assessment and status report on the Caribou (*Rangifer tarandus*), Newfoundland population, Atlantic-Gaspésie population and Boreal population, in Canada. Comnmittee on the Status of Endangered Wildlife in Canada. Available from (www.registrelep-sararegistry.gc.ca/default_e.cfm.

11. Courchamp, F., Clutton-Brock, T., and Grenfell, B. 1999. Inverse density dependence and the Allee effect. Trends in Ecology & Evolution 14(10): 405–410. doi:10.1016/S0169-5347(99)01683-3.

12. Courtois, R., Gingras, A., Fortin, D., Sebbane, A., Rochette, B., and Breton, L. 2008. Demographic and behavioural response of woodland caribou to forest harvesting. Can. J. For. Res. 38(11): 2837– 2849. NRC Research Press. doi:10.1139/X08-119.

13. Creel, S., Schuette, P., and Christianson, D. 2014. Effects of predation risk on group size, vigilance, and foraging behavior in an African ungulate community. Behavioral Ecology 25(4): 773–784. doi:10.1093/beheco/aru050.

14. Cumming, H.G., Beange, D.B., and Lavoie, G. 1996. Habitat partitioning between woodland caribou and moose in Ontario: the potential role of shared prédation risk. Rangifer: 81–94. doi:10.7557/2.16.4.1224.

15. Dekelaita, D., Krausman, P., and Mahoney, S. 2022. Estimated effects of clear-cuts and burns associated with habitat use by female Newfoundland Caribou (*Rangifer tarandus*). The Canadian Field-Naturalist 136(4): 316–332. doi:10.22621/cfn.v136i4.2767.

16. DeMars, C.A., and Boutin, S. 2018. Nowhere to hide: Effects of linear features on predator–prey dynamics in a large mammal system. Journal of Animal Ecology 87(1): 274–284. doi:10.1111/1365-2656.12760.

17. DeMars, C.A., Serrouya, R., Mumma, M.A., Gillingham, M.P., McNay, R.S., and Boutin, S. 2019. Moose, caribou, and fire: have we got it right yet? Canadian Journal of Zoology 97(10): 866–879. doi:10.1139/cjz-2018-0319.

18. Dickie, M., McNay, S.R., Sutherland, G.D., Cody, M., and Avgar, T. 2020. Corridors or risk? Movement along, and use of, linear features varies predictably among large mammal predator and prey species. Journal of Animal Ecology 89(2): 623–634. doi:10.1111/1365-2656.13130.

19. Dickie, M., Serrouya, R., McNay, R.S., and Boutin, S. 2017. Faster and farther: wolf movement on linear features and implications for hunting behaviour. Journal of Applied Ecology 54(1): 253–263. doi:10.1111/1365-2664.12732.

20. Dyer, S.J., O’Neill, J.P., Wasel, S.M., and Boutin, S. 2001. Avoidance of industrial development by woodland caribou. The Journal of Wildlife Management 65(3): 531–542. doi:10.2307/3803106.

21. Dyer, S.J., O’Neill, J.P., Wasel, S.M., and Boutin, S. 2002. Quantifying barrier effects of roads and seismic lines on movements of female woodland caribou in northeastern Alberta. Canadian Journal of Zoology 80(5): 839–845. doi:10.1139/z02-060.

22. Eacker, D.R., Jakes, A.F., and Jones, P.F. 2023. Spatiotemporal risk factors predict landscape-scale survivorship for a northern ungulate. Ecosphere 14(2): e4341. doi:10.1002/ecs2.4341.

23. Festa-Bianchet, M., Ray, J.C., Boutin, S., Côté, S.D., and Gunn, A. 2011. Conservation of caribou (*Rangifer tarandus*) in Canada: an uncertain future. Canadian Journal of Zoology 89(5): 419–434. doi:10.1139/z11-025.

24. Hart, S.A., and Chen, H.Y.H. 2006. Understory vegetation dynamics of North American boreal forests. Critical Reviews in Plant Sciences 25(4): 381–397. doi:10.1080/07352680600819286.

25. He, P., Maldonado-Chaparro, A.A., and Farine, D.R. 2019. The role of habitat configuration in shaping social structure: a gap in studies of animal social complexity. Behavioral Ecology and Sociobiology 73(9): 1–14.

26. Heeres, R.W., Leclerc, M., Frank, S., Kopatz, A., Pelletier, F., and Zedrosser, A. 2024. Are nonsocial species more social than we think? Seasonal patterns in sociality in a solitary terrestrial carnivore. Animal Behaviour 216: 107–130. doi:10.1016/j.anbehav.2024.07.022.

27. Hendrix, J.G., Robitaille, A.L., Kusch, J.M., Webber, Q.M.R., and Vander Wal, E. 2024. Faithful pals and familiar locales: differentiating social and spatial site fidelity during reproduction. Philosophical Transactions of the Royal Society B: Biological Sciences 379(1912): 20220525. doi:10.1098/rstb.2022.0525.

28. Hermosilla, T., Wulder, M.A., White, J.C., and Coops, N.C. 2022. Land cover classification in an era of big and open data: Optimizing localized implementation and training data selection to improve mapping outcomes. Remote Sensing of Environment 268: 112780. doi:10.1016/j.rse.2021.112780.

29. Hornseth, M.L., and Rempel, R.S. 2016. Seasonal resource selection of woodland caribou (*Rangifer tarandus caribou*) across a gradient of anthropogenic disturbance. Canadian Journal of Zoology 94(2): 79–93. doi:10.1139/cjz-2015-0101.

30. Joanisse, G.D., Bradley, R.L., Preston, C.M., and Munson, A.D. 2007. Soil enzyme inhibition by condensed litter tannins may drive ecosystem structure and processes: the case of *Kalmia angustifolia*. New Phytologist 175(3): 535–546. doi:10.1111/j.1469-8137.2007.02113.x.

31. Johnson, C.J., Parker, K.L., Heard, D.C., and Seip, D.S. 2004. Movements, foraging habits, and habitat use strategies of northern woodland caribou during winter: Implications for forest practices in British Columbia. Journal of Ecosystems and Management. doi:10.22230/jem.2004v5n1a290.

32. Johnson, E. a., Miyanishi, K., and Weir, J.M.H. 1998. Wildfires in the western Canadian boreal forest: Landscape patterns and ecosystem management. Journal of Vegetation Science 9(4): 603–610. doi:10.2307/3237276.

33. Joly, K., Dale, B.W., Collins, W.B., and Adams, L.G. 2003. Winter habitat use by female caribou in relation to wildland fires in interior Alaska. Can. J. Zool. 81(7): 1192–1201. doi:10.1139/z03-109.

34. Kirchmeier-Young, M.C., Gillett, N.P., Zwiers, F.W., Cannon, A.J., and Anslow, F.S. 2019. Attribution of the influence of human-induced climate change on an extreme fire season. Earth’s Future 7(1): 2– 10. doi:10.1029/2018EF001050.

35. Konkolics, S., Dickie, M., Serrouya, R., Hervieux, D., and Boutin, S. 2021. A burning question: What are the implications of forest fires for woodland caribou? The Journal of Wildlife Management 85(8): 1685–1698. doi:10.1002/jwmg.22111.

36. Lafontaine, A., Drapeau, P., Fortin, D., Gauthier, S., Boulanger, Y., and St-Laurent, M.-H. 2019. Exposure to historical burn rates shapes the response of boreal caribou to timber harvesting. Ecosphere 10(5): e02739. doi:10.1002/ecs2.2739.

37. Lake, N.F., Arsenault, A., and Cwynar, L. 2024. A Holocene fire history from Terra Nova National Park, Newfoundland, Canada: vegetation and climate change both influenced the fire regime. Front. Ecol. Evol. 12. doi:10.3389/fevo.2024.1419121.

38. Landau, W.M. 2021. The targets R package: a dynamic Make-like function-oriented pipeline toolkit for reproducibility and high-performance computing. Journal of Open Source Software 6(57): 2959. doi:10.21105/joss.02959.

39. Latham, A.D.M., Latham, M.C., Mccutchen, N.A., and Boutin, S. 2011. Invading white-tailed deer change wolf–caribou dynamics in northeastern Alberta. The Journal of Wildlife Management 75(1): 204–212. doi:10.1002/jwmg.28.

40. Le Goff, M., Hendrix, J.G., Webber, Q.M.R., Robitaille, A.L., and Vander Wal, E. 2024. Environmental, social and morphological drivers of fission–fusion dynamics in a social ungulate. Animal Behaviour 207: 267–276. doi:10.1016/j.anbehav.2023.10.014.

41. Leblond, M., Dussault, C., and Ouellet, J. –P. 2013. Avoidance of roads by large herbivores and its relation to disturbance intensity. Journal of Zoology 289(1): 32–40. doi:10.1111/j.1469-7998.2012.00959.x.

42. Leblond, M., Frair, J., Fortin, D., Dussault, C., Ouellet, J.-P., and Courtois, R. 2011. Assessing the influence of resource covariates at multiple spatial scales: an application to forest-dwelling caribou faced with intensive human activity. Landscape Ecol 26(10): 1433–1446. doi:10.1007/s10980-011-9647-6.

43. Leclerc, M., Dussault, C., and St-Laurent, M.-H. 2012. Multiscale assessment of the impacts of roads and cutovers on calving site selection in woodland caribou. Forest Ecology and Management 286: 59– 65. doi:10.1016/j.foreco.2012.09.010.

44. Lesmerises, F., Déry, F., Johnson, C.J., and St-Laurent, M.-H. 2018a. Spatiotemporal response of mountain caribou to the intensity of backcountry skiing. Biological Conservation 217: 149–156. doi:10.1016/j.biocon.2017.10.030.

45. Lesmerises, F., Johnson, C.J., and St-Laurent, M.-H. 2018b. Landscape knowledge is an important driver of the fission dynamics of an alpine ungulate. Animal Behaviour 140: 39–47. doi:10.1016/j.anbehav.2018.03.014.

46. Mahoney, S.P., Lewis, K.P., Weir, J.N., Morrison, S.F., Glenn Luther, J., Schaefer, J.A., Pouliot, D., and Latifovic, R. 2016. Woodland caribou calf mortality in Newfoundland: insights into the role of climate, predation and population density over three decades of study. Population Ecology 58(1): 91–103. doi:10.1007/s10144-015-0525-y.

47. Makuya, L., and Schradin, C. 2024. The secret social life of solitary mammals. Proceedings of the National Academy of Sciences 121(13): e2402871121. doi:10.1073/pnas.2402871121.

48. Mallik, A., and Kayes, I. 2018. Lichen matted seedbeds inhibit while moss dominated seedbeds facilitate black spruce (*Picea mariana*) seedling regeneration in post-fire boreal forest. Forest Ecology and Management 427: 260–274. doi:10.1016/j.foreco.2018.05.064.

49. Mallik, A.U. 1995. Conversion of temperate forests into heaths: Role of ecosystem disturbance and ericaceous plants. Environmental Management 19(5): 675–684. doi:10.1007/BF02471950.

50. Mallik, A.U., Bloom, R.G., and Whisenant, S.G. 2010. Seedbed filter controls post-fire succession. Basic and Applied Ecology 11(2): 170–181. doi:10.1016/j.baae.2009.11.005.

51. Maltman, J.C., Coops, N.C., Rickbeil, G.J.M., Hermosilla, T., and Burton, A.C. 2024. Quantifying forest disturbance regimes within caribou (*Rangifer tarandus*) range in British Columbia. Sci Rep 14(1): 6520. doi:10.1038/s41598-024-56943-0.

52. Marsh, C.D., and Clawson, A.B. 1930. Mountain laurel (*Kalmia latifolia*) and sheep laurel (*Kalmia angustifolia*) as stock-poisoning plants. Technical Bulletin, US Department of Agriculture.

53. Matthiopoulos, J., Fieberg, J.R., and Aarts, G. 2023. Species-Habitat Associations: Spatial data, predictive models, and ecological insights, 2nd Edition. University of Minnesota Libraries Publishing. Available from https://hdl.handle.net/11299/217469 [accessed 19 September 2024].

54. McCarthy, S.C., Weladji, R.B., Doucet, C., and Saunders, P. 2011. Woodland caribou calf recruitment in relation to calving/post-calving landscape composition. Rangifer 31(1): 35–47. doi:10.7557/2.31.1.1918.

55. McNamara, J.A., Schaefer, J.A., Bastille-Rousseau, G., and Mahoney, S.P. 2022. Landscape features and caribou harvesting during three decades in Newfoundland. Ecoscience 29(1): 39–53. Taylor & Francis. doi:10.1080/11956860.2021.1969825.

56. Merkle, J.A., Poulin, M.-P., Caldwell, M.R., Laforge, M.P., Scholle, A.E., Verzuh, T.L., and Geremia, C. 2024. Spatial–social familiarity complements the spatial–social interface: evidence from Yellowstone bison. Philosophical Transactions of the Royal Society B: Biological Sciences 379(1912): 20220530. doi:10.1098/rstb.2022.0530.

57. Mooring, M.S., Fitzpatrick, T.A., Nishihira, T.T., and Reisig, D.D. 2004. Vigilance, predation risk, and the Allee effect in desert bighorn sheep. The Journal of Wildlife Management 68(3): 519–532. doi:10.2193/0022-541X(2004)068[0519:VPRATA]2.0.CO;2.

58. Mumma, M.A., Gillingham, M.P., Parker, K.L., Johnson, C.J., and Watters, M. 2018. Predation risk for boreal woodland caribou in human-modified landscapes: Evidence of wolf spatial responses independent of apparent competition. Biological Conservation 228: 215–223. doi:10.1016/j.biocon.2018.09.015.

59. Nagy-Reis, M., Dickie, M., Calvert, A.M., Hebblewhite, M., Hervieux, D., Seip, D.R., Gilbert, S.L., Venter, O., DeMars, C., Boutin, S., and Serrouya, R. 2021. Habitat loss accelerates for the endangered woodland caribou in western Canada. Conservation Science and Practice 3(7): e437. doi:10.1111/csp2.437.

60. Padgham, M., Lovelace, R., Salmon, M., and Rudis, B. 2017. osmdata. Journal of Open Source Software 2(14): 305. doi:10.21105/joss.00305.

61. Parks Canada Agency. (n.d.). Fire Management Plan for Terra Nova National Park, 2022-2032.

62. Pebesma, E. 2018. Simple Features for R: Standardized Support for Spatial Vector Data. The R Journal10(1): 439–446.

63. Peignier, M., Webber, Q.M.R., Robitaille, A.L., Koen, E.L., Laforge, M.P., and Wal, E.V. 2019. Space use and social association in a gregarious ungulate: testing the conspecific attraction and resource dispersion hypotheses. Ecology and Evolution: 1–13. doi:10.1002/ece3.5071.

64. Prokopenko, C.M., Ellington, E.H., Robitaille, A., Aubin, J.A., Balluffi-Fry, J., Laforge, M., Webber, Q.M.R., Zabihi-Seissan, S., and Vander Wal, E. 2024. Friends because of foes: synchronous movement within predator–prey domains. Philosophical Transactions of the Royal Society B: Biological Sciences 379(1912): 20230374. doi:10.1098/rstb.2023.0374.

65. Rayl, N.D., Fuller, T.K., Organ, J.F., McDonald, J.E., Jr., Mahoney, S.P., Soulliere, C., Gullage, S.E., Hodder, T., Norman, F., Porter, T., Bastille-Rousseau, G., Schaefer, J.A., and Murray, D.L. 2014. Mapping the distribution of a prey resource: neonate caribou in Newfoundland. Journal of Mammalogy 95(2): 328–339. doi:10.1644/13-MAMM-A-133.1.

66. Rettie, W.J., and Messier, F. 2000. Hierarchical habitat selection by woodland caribou: Its relationship to limiting factors. Ecography 23(4): 466–478. doi:10.1111/j.1600-0587.2000.tb00303.x.

67. Rickbeil, G.J.M., Hermosilla, T., Coops, N.C., White, J.C., and Wulder, M.A. 2017. Barren-ground caribou (*Rangifer tarandus groenlandicus*) behaviour after recent fire events; integrating caribou telemetry data with Landsat fire detection techniques. Global Change Biology 23(3): 1036–1047. doi:10.1111/gcb.13456.

68. Robitaille, A.L., Webber, Q.M.R., and Vanderwal, E. 2019. Conducting social network analysis with animal telemetry data: Applications and methods using spatsoc. Methods in Ecology and Evolution 00: 1–9. doi:10.1111/2041-210X.13215.

69. Russell, K.L.M. 2018. Close encounters of the burned kind: Spatiotemporal effects of fire on habitat selection strategies of woodland caribou (*Rangifer tarandus caribou*) during winter. MSc, University of Northern British Columbia. doi:10.24124/2018/58873.

70. Schindler, D.W., Walker, D., Davis, T., and Westwood, R. 2007. Determining effects of an all weather logging road on winter woodland caribou habitat use in south-eastern Manitoba. Rangifer: 209– 217. doi:10.7557/2.27.4.346.

71. Serrouya, R., Dickie, M., DeMars, C., Wittmann, M.J., and Boutin, S. 2020. Predicting the effects of restoring linear features on woodland caribou populations. Ecological Modelling 416: 108891. doi:10.1016/j.ecolmodel.2019.108891.

72. Signer, J., Fieberg, J., and Avgar, T. 2019. Animal movement tools (amt): R package for managing tracking data and conducting habitat selection analyses. Ecology and Evolution 9(2): 880–890. doi:10.1002/ece3.4823.

73. Silk, J.B. 2007. The adaptive value of sociality in mammalian groups. Philosophical Transactions of the Royal Society B: Biological Sciences 362(1480): 539–559. Royal Society. doi:10.1098/rstb.2006.1994.

74. Silva, J.A., Nielsen, S.E., McLoughlin, P.D., Rodgers, A.R., Hague, C., and Boutin, S. 2020. Comparison of pre-fire and post-fire space use reveals varied responses by woodland caribou (*Rangifer tarandus caribou*) in the Boreal Shield. Canadian Journal of Zoology 98(11): 751–760. doi:10.1139/cjz-2020-0139.

75. Skatter, H.G., Charlebois, M.L., Eftestøl, S., Tsegaye, D., Colman, J.E., Kansas, J.L., Flydal, K., and Balicki, B. 2017. Living in a burned landscape: woodland caribou (*Rangifer tarandus caribou*) use of postfire residual patches for calving in a high fire – low anthropogenic Boreal Shield ecozone. Can. J. Zool. 95(12): 975–984. doi:10.1139/cjz-2016-0307.

76. Smith, A., and Johnson, C.J. 2023. Why didn’t the caribou (*Rangifer tarandus groenlandicus*) cross the winter road? The effect of industrial traffic on the road-crossing decisions of caribou. Biodiversity and Conservation 32(8): 2943–2959. doi:10.1007/s10531-023-02637-4.

77. Sorensen, T., McLoughlin, P.D., Hervieux, D., Dzus, E., Nolan, J., Wynes, B., and Boutin, S. 2008. Determining sustainable levels of cumulative effects for boreal caribou. Journal of Wildlife Management 72(4): 900–905. doi:10.2193/2007-079.

78. Stevens-Rumann, C.S., Prichard, S.J., Whitman, E., Parisien, M.-A., and Meddens, A.J.H. 2022.Considering regeneration failure in the context of changing climate and disturbance regimes in western North America. Canadian Journal of Forest Research 52(10): 1281–1302. doi:10.1139/cjfr-2022-0054.

79. Stewart, F.E.C., Nowak, J.J., Micheletti, T., McIntire, E.J.B., Schmiegelow, F.K.A., and Cumming, S.G. 2020. Boreal caribou can coexist with natural but not industrial disturbances. The Journal of Wildlife Management 84(8): 1435–1444. doi:10.1002/jwmg.21937.

80. Uccheddu, S., Body, G., Weladji, R.B., Holand, Ø., and Nieminen, M. 2015. Foraging competition in larger groups overrides harassment avoidance benefits in female reindeer (*Rangifer tarandus*). Oecologia 179(3): 711–718. doi:10.1007/s00442-015-3392-5.

81. Viana, D.S., Granados, J.E., Fandos, P., Pérez, J.M., Cano-Manuel, F.J., Burón, D., Fandos, G., Aguado, M.Á.P., Figuerola, J., and Soriguer, R.C. 2018. Linking seasonal home range size with habitat selection and movement in a mountain ungulate. Movement Ecology 6(1): 1. doi:10.1186/s40462-017-0119-8.

82. Viejou, R., Avgar, T., Brown, G.S., Patterson, B.R., Reid, D.E.B., Rodgers, A.R., Shuter, J., Thompson, I.D., and Fryxell, J.M. 2018. Woodland caribou habitat selection patterns in relation to predation risk and forage abundance depend on reproductive state. Ecology and Evolution 8(11): 5863– 5872. doi:10.1002/ece3.4124.

83. Walker, P.D., Rodgers, A.R., Shuter, J., Fryxell, J.M., and Merrill, E.H. 2024. Woodland caribou calving fidelity: Spatial location, habitat, or both? Ecology and Evolution 14(6): e11480. doi:10.1002/ece3.11480.

84. Webber, Q., Prokopenko, C., Kingdon, K., Turner, J., and Vander Wal, E. 2024a. Effects of the social environment on movement-integrated habitat selection. Movement Ecology 12(1): 61. doi:10.1186/s40462-024-00502-9.

85. Webber, Q.M.R., Ferraro, K., Hendrix, J., and Vander Wal, E. 2022. What do caribou eat? A review of the literature on caribou diet. Canadian Journal of Zoology. doi:10.1139/cjz-2021-0162.

86. Webber, Q.M.R., Laforge, M.P., Bonar, M., and Vander Wal, E. 2024b. The adaptive value of density-dependent habitat specialization and social network centrality. Nature Communications 15(1): 4423. doi:10.1038/s41467-024-48657-8.

87. Webber, Q.M.R., and Vander Wal, E. 2018. An evolutionary framework outlining the integration of individual social and spatial ecology. Journal of Animal Ecology 87(1): 113–127. doi:10.1111/1365-2656.12773.

88. Webber, Q.M.R., and Vander Wal, E. 2021. Context-dependent group size: effects of population density, habitat, and season. Behavioral Ecology 32(5): 970–981. doi:10.1093/beheco/arab070.

89. Wilson, R.R., Parrett, L.S., Joly, K., and Dau, J.R. 2016. Effects of roads on individual caribou movements during migration. Biological Conservation 195: 2–8. doi:10.1016/j.biocon.2015.12.035.

90. Wittmer, H.U., Sinclair, A.R.E., and McLellan, B.N. 2005. The role of predation in the decline and extirpation of woodland caribou. Oecologia 144(2): 257–267. doi:10.1007/s00442-005-0055-y.

