## Appendix for "Social facilitation of risky habitats in woodland caribou: responses to fire and roads"

Supplementary Materials

Highlight: Assessment and protection of preferred habitat based on behaviour during very low density may not capture the breadth of habitats used by a healthy population.

*Road model*

This same model structure was first applied to all data, and then independently to each of the four biological seasons.

Step (observed/available) ~ log(step length) +

log(step length) : forest +

forest +

log(step length) : open +

open +

log(distance to TCH +1) +

log(distance to TCH +1) : log(step length) +

log(distance to minor roads +1) +

log(distance to minor roads +1) : log(step length) +

(1 | step ID) +

log(step length) | animal ID +

log(step length) : forest | animal ID +

forest | animal ID +

open | animal ID +

log(distance to TCH +1) | animal ID +

log(distance to TCH +1) : log(step length) | animal ID +

log(distance to minor roads +1) | animal ID +

log(distance to minor roads +1) : log(step length) | animal ID

*Fire model*

This same model structure was first applied to all data, and then independently to each of the four biological seasons.

Step (observed/available) ~ log(step length) +

log(step length) : forest +

forest +

log(step length) : open +

open +

log(distance to new burn +1) +

log(distance to new burn +1) : log(step length) +

log(distance to old burn +1) +

log(distance to old burn +1) : log(step length) +

(1 | step ID) +

log(step length) | animal ID +

log(step length) : forest | animal ID +

forest | animal ID +

open | animal ID +

log(distance to new burn +1) | animal ID +

log(distance to new burn +1) : log(step length) | animal ID +

log(distance to old burn +1) | animal ID +

log(distance to old burn +1) : log(step length) | animal ID

*Social road model*

Applied only to winter data (additional terms relative to base road model are **bolded**).

Step (observed/available) ~ log(step length) +

log(step length) : forest +

forest +

**forest : in dyad +**

log(step length) : open +

open +

**open : in dyad +**

log(distance to TCH +1) +

**log(distance to TCH +1) : in dyad +**

log(distance to TCH +1) : log(step length) +

log(distance to minor roads +1) +

**log(distance to minor roads +1) : in dyad +**

log(distance to minor roads +1) : log(step length) +

(1 | step ID) +

log(step length) | animal ID +

log(step length) : forest | animal ID +

forest | animal ID +

open | animal ID +

log(distance to TCH +1) | animal ID +

log(distance to TCH +1) : log(step length) | animal ID +

log(distance to minor roads +1) | animal ID +

log(distance to minor roads +1) : log(step length) | animal ID

*Social fire model*

Applied only to winter data (additional terms relative to base fire model are **bolded**).

Step (observed/available) ~ log(step length) +

log(step length) : forest +

forest +

**forest : in dyad +**

log(step length) : open +

open +

**open : in dyad +**

log(distance to new burn +1) +

**log(distance to new burn +1) : in dyad +**

log(distance to new burn +1) : log(step length) +

log(distance to old burn +1) +

**log(distance to old burn +1) : in dyad +**

log(distance to old burn +1) : log(step length) +

(1 | step ID) +

log(step length) | animal ID +

log(step length) : forest | animal ID +

forest | animal ID +

open | animal ID +

log(distance to new burn +1) | animal ID +

log(distance to new burn +1) : log(step length) | animal ID +

log(distance to old burn +1) | animal ID +

log(distance to old burn +1) : log(step length) | animal ID

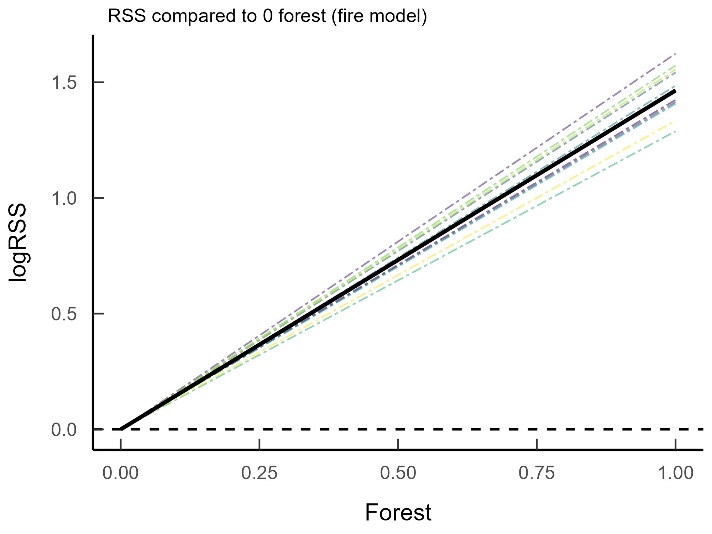

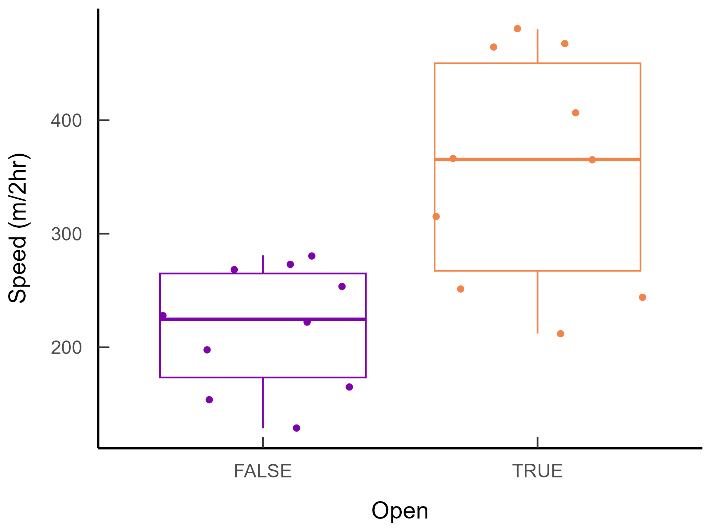

Figure S1. (A) Relative selection strength (RSS) for forest habitats. Each coloured dashed line represents an individual caribou, with the solid black line showing the population average. RSS is the strength of selection for a given value of a covariate, compared to a reference level. In this case values between 0 and 100% forest are compared against a reference level of 0 forest. Note the y-axis is log() transformed: for example, given the choice between 75% forest and 0 forest, caribou are 10 times more likely to select the 75% forest location (when log(RSS) = 1, RSS = 10). (B) Effect of habitat openness on movement rate of caribou. Each point represents an individual caribou, with boxplots showing the population overall. After accounting for other covariates, predicted step lengths between consecutive 2 hr fixes are longer in open habitats than in non-open areas. Caribou thus move faster in open areas in general.

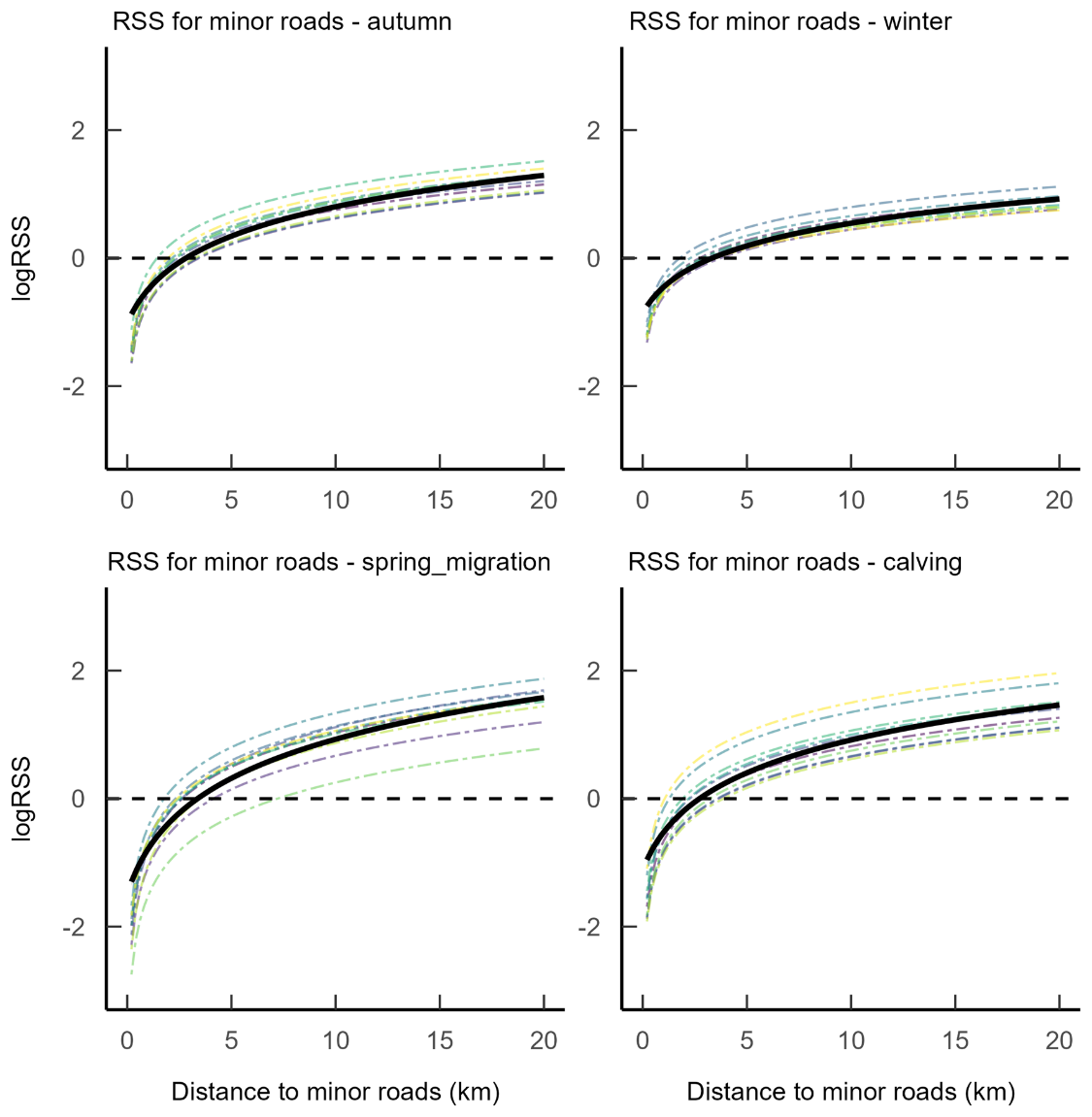

Figure S2. Seasonal differences in relative selection strength (RSS) of caribou in response to minor roads in and around Terra Nova National Park. Each coloured dashed line represents an individual caribou, with the solid black line showing the population average. RSS is the strength of selection for a given value of a covariate, compared to a reference level. In this case values between 0 and 20 km from minor roads are compared against a reference level of the median distance to minor roads (approx. 4 km). Caribou avoided minor roads during all seasons, with minimal variation throughout the year.

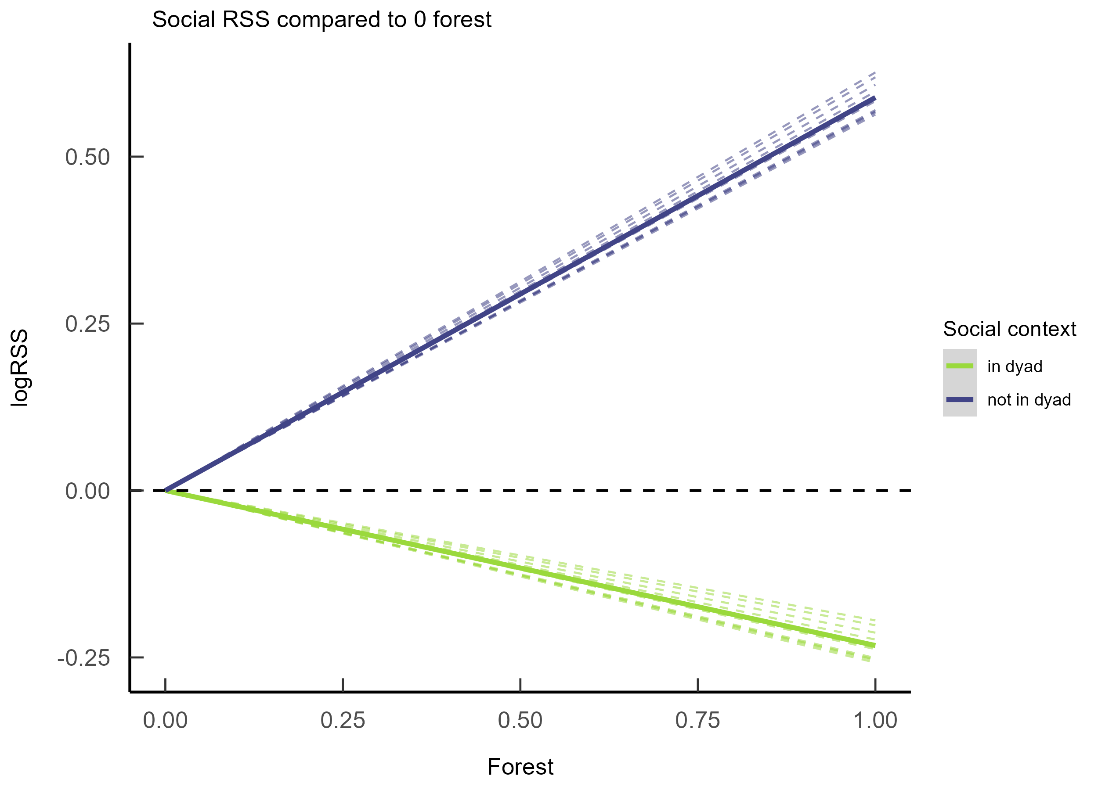

Figure S3. Relative selection strength (RSS) for forested habitats of caribou in dyads with other collared individuals vs. when not in a dyad, during winter. Green lines represent periods when caribou were in a dyad with other collared individuals. Blue lines are periods when no other collared individuals were within 50m of the focal animal. RSS is the strength of selection for a given value of a covariate, compared to a reference level. In this case values between 0 and 100% forest are compared against a reference level of 0 forest. Caribou selected for forest when not in a dyad, but when in a dyad with conspecifics, forests were avoided.
